# Social network centrality predicts dietary decisions in a wild bird population

**DOI:** 10.1101/2023.08.25.554636

**Authors:** Keith McMahon, Nicola M. Marples, Lewis G. Spurgin, Hannah M. Rowland, Ben C. Sheldon, Josh A. Firth

## Abstract

How individuals balance costs and benefits of group living remains central to understanding sociality. In relation to diet, social foraging provides many advantages but also increases competition. Nevertheless, social individuals may offset increased competition by broadening their diet and consuming novel foods. Despite the expected relationships between social behaviour and dietary decisions, how sociality shapes individuals’ novel food consumption remains largely untested in natural populations. Here, we use wild, RFID-tracked, great tits to experimentally test how sociality predicts dietary decisions. We show that individuals with more social connections have higher propensity to use novel foods compared to socially-peripheral individuals, and this is unrelated to neophobia, observations, and demographic factors. These findings indicate sociable individuals may offset potential costs of competition by foraging more broadly. We discuss how social environments may drive behavioural change in natural populations, and the implications for the causes and consequences of social strategies and dietary decisions.

## INTRODUCTION

Groups of foraging animals benefit from sociality in many ways ^1^, ranging from social foraging allowing complex cooperative hunting ^2^ to other benefits such as learning about how food is distributed or avoiding predators ^1–5^. It is also likely that social learning may help individuals decide which new foods to explore and consume ^6–8^. Indeed, novel foods pose a challenge to foragers because they can differ from familiar foods in their nutritional quality, and may also contain unfavorable chemicals or defensive toxins ^9^. How individual foragers in the wild differ in their propensity to explore and use novel foods and how individual sociality affects foraging decisions about novel foods remains a central topic in understanding dietary decision making [10].

As well as many benefits, foraging in groups also imposes costs through intraspecific competition ^10^. When resources are limited, there is greater competition between group members ^11^ and this can result in carry-over costs through ‘interference competition’ e.g. fighting for resources, with attendant costs in time and risk of injury ^12^. This interference competition can reduce the time available to an individual to make decisions about which food items to consume, and can reduce the number of profitable food items encountered while foraging ^13^. Exploiting new food sources, in this situation, can alleviate within-group competition through niche partitioning ^14–16^.

When faced with decisions about which new foods to explore and consume, many species exhibit dietary wariness ^17,18^. Dietary wariness is composed of two behavioural processes: neophobia and dietary conservatism ^19^. Neophobia describes the initial fear/apprehension of novel objects or foods, and is observed in many animal groups, including fish, mammals and birds ^18,20,21^. This aversion is usually brief, and is followed by investigation of the novel food or object ^22^. Once neophobia has waned, some individuals within a population continue to avoid novel food long after the initial exposure, and this is termed dietary conservatism ^19^. Dietary conservatism is a spectrum: individuals differ in their willingness to consume novel food. Adventurous consumers are those individuals which show little or no hesitation in consuming novel food once neophobia has passed, while conservatively foraging individuals continue to avoid the novel food for extended periods ^23^. Following this general classification, dietary conservatism has been observed in a wide variety of species, particularly in various studies in birds ^21,24^ and fish ^25–27^. It is important to consider individual differences in the propensity to eat novel foods when discussing the strategies that animals use to mitigate resource competition during social foraging, as this is of direct relevance for the study of the factors that shape sociality and resource acquisition ^28^ .

The advent of animal tracking technologies has revolutionised our ability to observe individuals’ social foraging associations in the wild ^29^, and animal social networks have now been quantified across a range of animal systems ^30^. Social network analysis provides a framework for quantifying variation in intraspecific sociality ^31^ and allows the estimation of various metrics of individuals’ social behaviour ^32^. This provides fine-scale information about an individual’s own social associations, as well as the wider social environment they inhabit ^33^, who they associate with, when they associate with them and where the associations happen. The ability to quantify social networks within the wild, while simultaneously tracking individuals’ foraging behaviour, presents the opportunity to determine empirically how intraspecific differences in sociality relate to the various aspects of dietary wariness in natural settings.

In this study, we use novel food experiments to test individual-level dietary wariness in a RFID-tracked social system of wild great tits (*Parus major)*. Using this approach, we are able to examine dietary wariness and novel food usage independently, and use social network analysis to determine how individual sociality predicts individuals’ foraging decisions. A priori we expected that there would be variation in the use of novels foods among foraging great tits. This variation would be driven in large part by underlying propensities to consume novel food, dietary wariness, but also that social network position would have a major role in influencing decisions made by foraging birds. Thus, we were able to directly test the expectation that more social individuals have a greater propensity to eat novel foods, whether that be due to the need to mitigate the potential costs of interference competition or because they have more access to information about the profitability of novel food sources. These potential explanations need not be mutually exclusive. As a consequence these more social individuals should show lower levels of dietary wariness compared to less social individuals. We are able to separate out other elements (observation-related factors and demographic traits) when assessing the relationship between foraging decisions and a suite of social measures. We discuss the implications of these experimental results for understanding how competition shapes social foraging, and the wider insights this may offer into the interplay between individual-foraging decisions and social behaviour.

## RESULTS

During the study, 105 unique RFID tagged great tits were detected: 85 during the baseline data collection period, and 75 and 61 in the first and second experimental trial, respectively. The average number of detections of each RFID tagged individual over the 19-day experiment was 3234±409 (mean±SE), with a total of 210,579 detections of all individuals during the baseline period and 60,727 and 68,311 during the first trial and second trial, respectively. We detected 2393 flocking events for the baseline period, and 767 and 764 for the first and second trail, respectively. The typical group size (i.e. group size encountered by the average individual ^34^ was 6.8±0.03. The social networks inferred from these flocking events (see Methods) were relatively dense networks within sites (Figure 1), with a total number of unweighted social network connections of 1266 in the baseline period and 892 and 697 for the first and second experimental trials, respectively.

**Figure 1.**
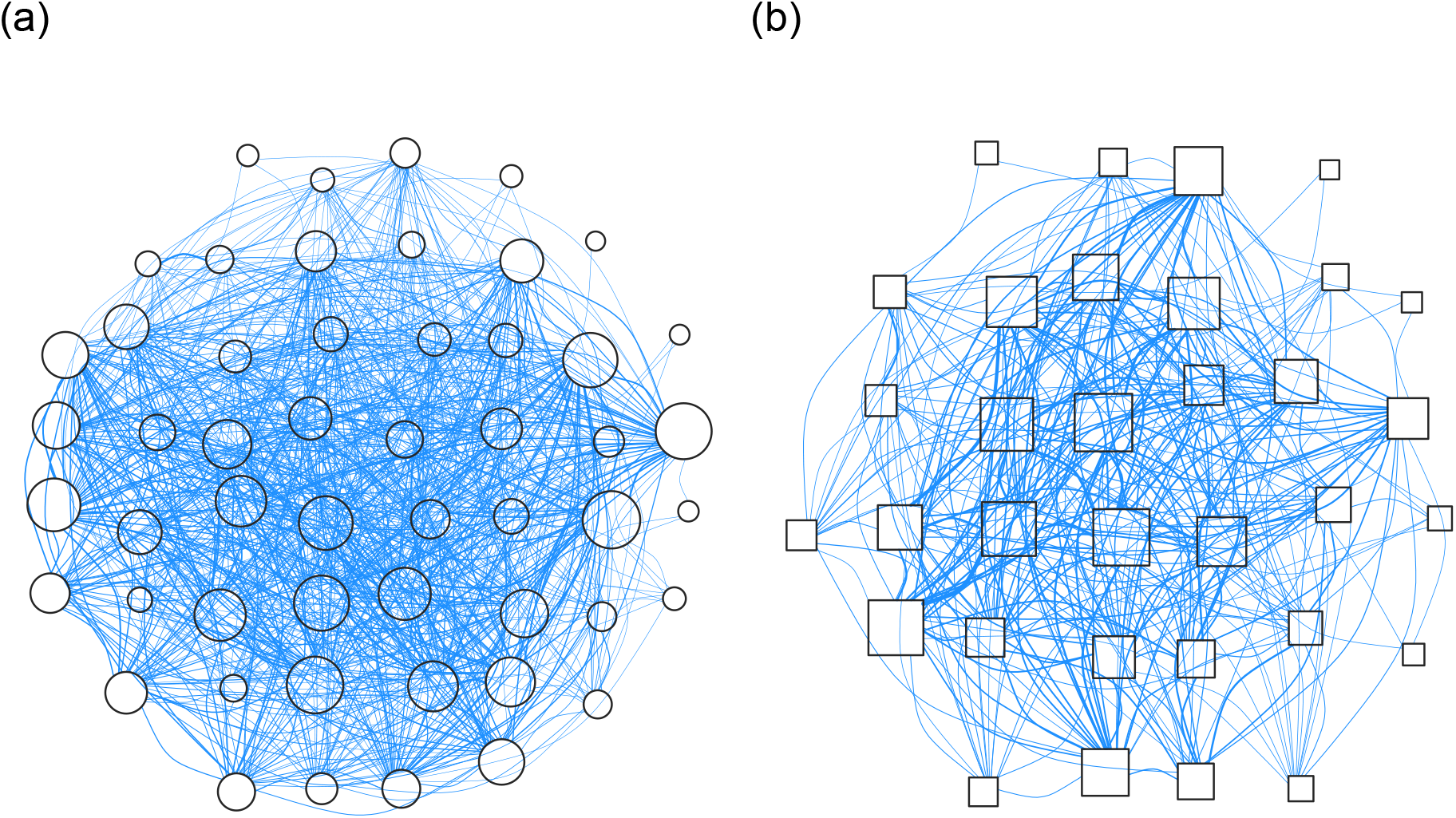
Social networks at the experimental sites. Site 1 (Figure 1a) and experimental site 2 (Figure 1b) in the baseline period. The nodes (points) represent the individuals and the edges (lines) show the social connections between them. The size of the nodes and their shading indicates an individual’s network centrality (large dark nodes = high strength, small light nodes = low strength) and are positioned using spring layout forced into a best-fit filled circle. The thickness of the lines shows the weighted social bond between dyads where thick lines indicate strongly connected individuals and thin lines show weak connections (the edge thickness is standardised by total sum of social connections with the network). Although site 1 (Figure 1a) appears to be denser than site 2 (Figure 1b), the actual network densities (percentage of potential realised links) are very similar (70% and 63% respectively), and the main visual difference comes from higher number of individuals in site 1 (nodes=52) resulting in more connections (connections=931) than site 2 (nodes=33, connections=335).

### Social Centrality and Novel Food Usage

An Individual’s propensity to use novel food during each of the experimental trials was significantly predicted by their prior social centrality (Figure 2): the GLMs showed a strong relationship between proportion of novel food usage and the individuals’ prior weighted strength for both trials (Trial 1 - Table S2a: Coefficient= 0.529±0.235, t=2.25, p=0.028, p_rand_=0.01. Trial 2 - Table S2b: Coef= 0.467±0.150, t=3.11, p=0.003, p_rand_=0.012). None of the other individual characteristics in the models (age, sex, immigrant status, previous feeder usage) were significant predictors of novel food usage (Table 2). The first experimental site had a strong colour preference for red over green when each colour was novel (Figure 2), the first trial had a reduced novel food usage for site 1 (initially using green novel food) over site 2 (using red novel food) and the reverse effect for the second trial when the novel food colours were swapped (Trial 1 - Table S2a: Coef= 3.40±0.71, t=4.8, p<0.001. Trial 2 - Table S2b: Coef= -1.59±0.27, t= -5.96, p<0.001). This apparent effect of colour preference persisted through all of the models (See Supplementary Tables).

**Figure 2.**
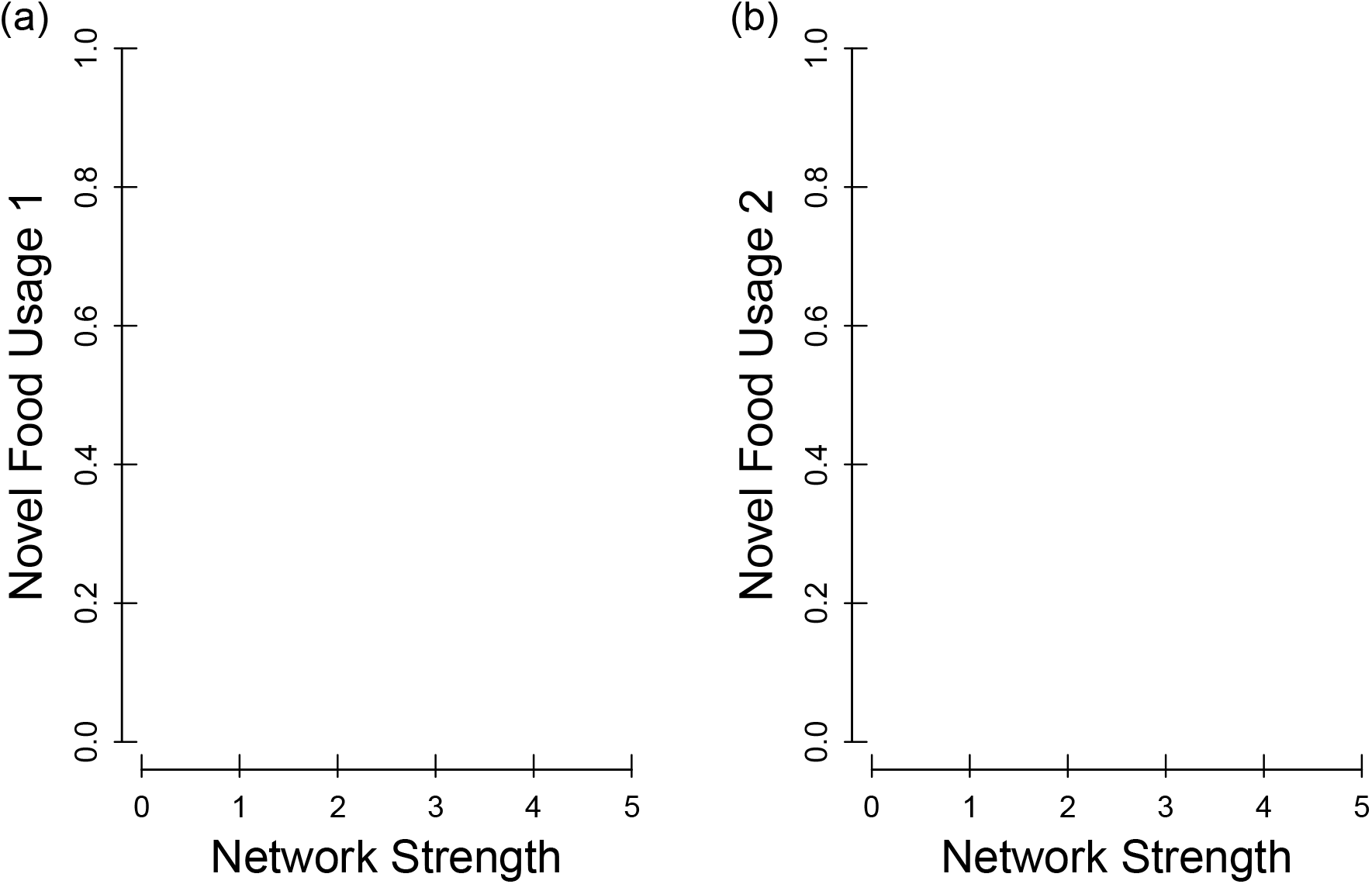
Social centrality and novel food use. Prior social centrality (network strength – x axis) and subsequent novel food usage (proportion of novel food usage – y axis) for the (a) first trial, and (b) second trial. The point positions show the individual data points, point colour shows the colour of the novel food (red or green dyed peanut), point shape shows which experimental site the individual was at (site 1 round, site 2 square), and point size indicates weight of the data point i.e. the total number of detections (at both the novel, and familiar food feeder). The lines show the GLM fit, and the surrounding polygons show the associated standard error around this estimate, with the red lines showing the fit for the red novel food site, the green line showing the fit for the green novel food site, and the black line denoting the overall fit. See Table S2 for full model details.

Supplementary analysis which considered two alternative measures of centrality (‘average edge weight’ and ‘eigenvector centrality’) confirmed the findings that prior social network position significantly predicted novel food usage. (Figure S1; Table S3-S4). The average edge weight was significantly related to the proportion of novel food usage across both trials (Trial 1 - Table S3a: Coef= 16.5±7.2, t=2.28, p=0.027, p_rand_=0.022. Trial 2 - Table S3b: Coef=15.9±4.5, t=3.5, p=0.001, p_rand_=0.004) as was eigenvector centrality (Trial 1 - Table S4a: Coef=1.74±0.76, t=2.29, p=0.026, p_rand_=0.05. Trial 2- Table S4b: Coef=1.71±0.58, t=2.93, p=0.005, p_rand_=0.012). A further line of supplementary analysis confirmed the importance of using network centrality as a robust measure of sociality, as novel food usage was not significantly related to more basic social measures (Figure S2; Table S5-S6) that simply quantified an individual’s average flock size (Trial 1 - Table S5a; Coef= 0.049±0.17, t=0.29, p=0.77, p_rand_=0.68. Trial 2 - Table S5b; Coef=-0.14±0.15, t=-0.92, p=0.36, p_rand_=0.23) or their total number of flock mates (Trial 1 - Table S6a; Coef=0.036±0.048, t=0.75, p=0.45, p_rand_=0.18. Trial 2 - Table S6b; Coef=0.0565±0.0385, t=1.47, p=0.15, p_rand_=0.10.)

### Novel Food Neophobia and Social Centrality

The majority of individuals (92%) recorded during the experimental trials were detected on the novel food feeder during the trial, indicating that complete neophobia (unwillingness to try the novel food at all) was extremely rare. Furthermore, 95% of those that were detected using the novel food feeder during the trial were recorded using it on the first day of the trial, again indicating that neophobia generally was not a persistent barrier to novel food usage.

However, we also aimed to examine whether any individual variation in initial avoidance of the novel food (i.e. neophobia) was related to individuals’ network position. By using the very first record of each bird during the experimental trial, we found that whether or not individuals perched on the novel food feeder hole when they first arrived at the experimental trial was not significantly related to social network centrality in either the first trial (Figure 3a – Coef=0.29±0.41, t=0.7, p=0.48, p_rand_=0.29, Table S7a) or the second trial (Figure 3b - Coef=-0.55±0.57, t=-0.96, p=0.34, p_rand_=0.28, Table S7b). Although only 30% of individuals immediately tried the novel food when first arriving at the experimental trials, none of the individual characteristics included in the GLM were predictive of which individuals perched on the novel food feeder hole as they first arrived during the experiment (Table S7).

**Figure 3.**
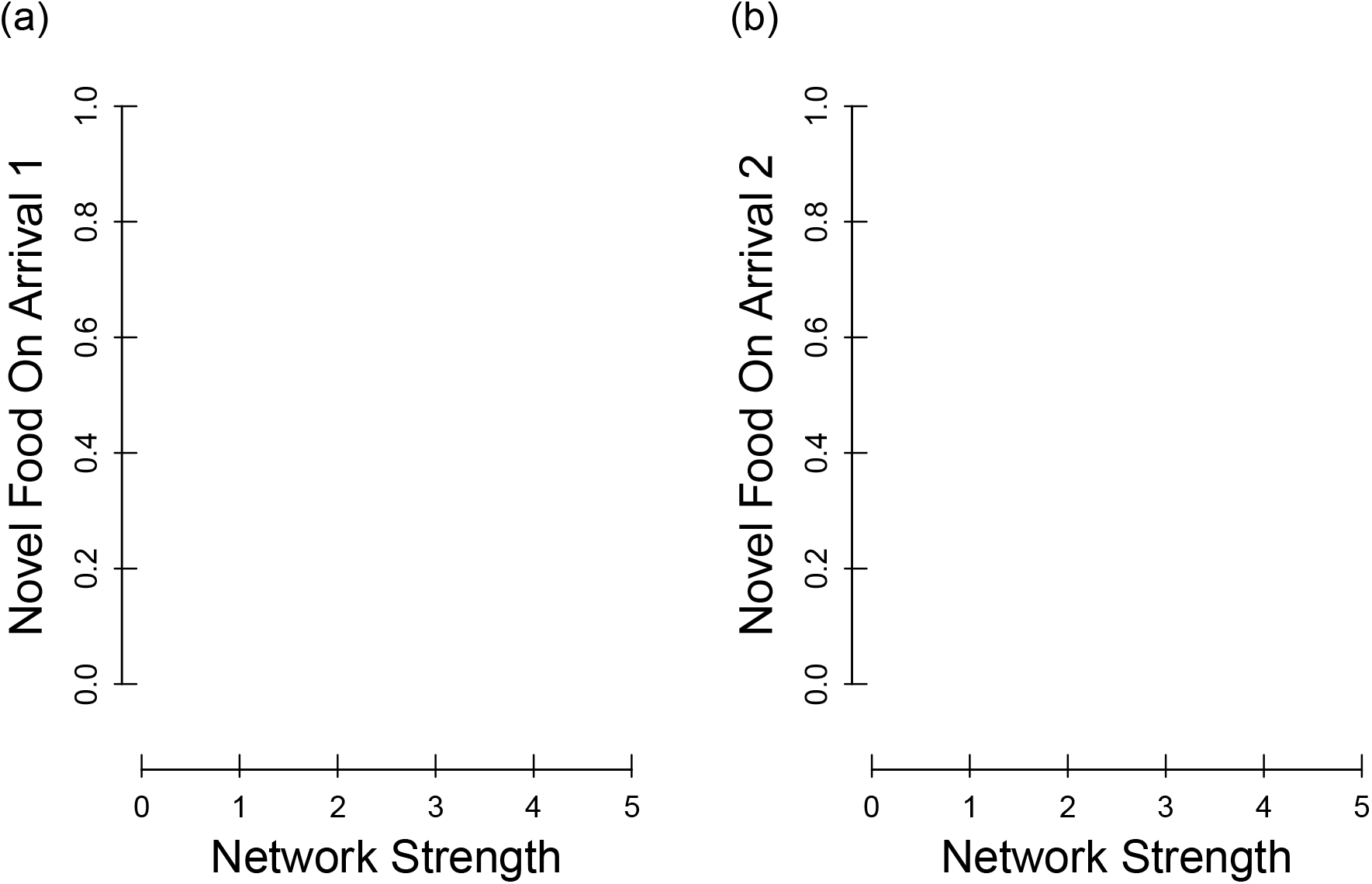
Social centrality and trying novel food on arrival. Social Prior social centrality (network strength – x axis) and probability of individuals trying the novel food upon arrival at the (a) first trial, and (b) second trial. The point positions show the individual network strength and whether they immediately tried the novel food (top) or not (bottom), point colour shows the colour of the novel food (red or green dyed peanut), and point shape shows which experimental site the individual was at (site 1 circles or site 2 squares). The lines show the GLM fit, and the surrounding polygons show the associated standard error around this estimate, with the red lines showing the fit for the red novel food site, the green line showing the fit for the green novel food site, and the black line denoting the overall fit. See Table S7 for full model details.

In line with this result, supplementary analysis also showed that network strength was not related to the amount of time taken for each individual to first land on the feeding perch of the novel food in each trial (Table S8;S9). This was true when time was quantified as the time of day they were first recorded on the novel food (Table S8), or when quantified as the total elapsed foraging time since they were first detected at the site during the trial (Table S9).

As a direct assessment of whether the relationship between sociality and proportional usage of novel food exists regardless of any neophobia, we also found that prior network strength significantly predicted the proportion of novel food (over familiar food) that individuals used after they had first tried the novel food feeder (Figure 4; Table S10) i.e. after any neophobia was overcome and only dietary conservatism was active. Again, this was true for both the first trial (Figure 4a - Coef=0.55±0.25, t=2.15, p=0.037,p_rand_=0.006, Table S10a) and second trial (Figure 4b - Coef=0.53±0.168, t=3.17, p=0.003,p_rand_=0.010, Table S10b), and the site/colour preference effect was again evident (Figure 4; Table S10).

**Figure 4.**
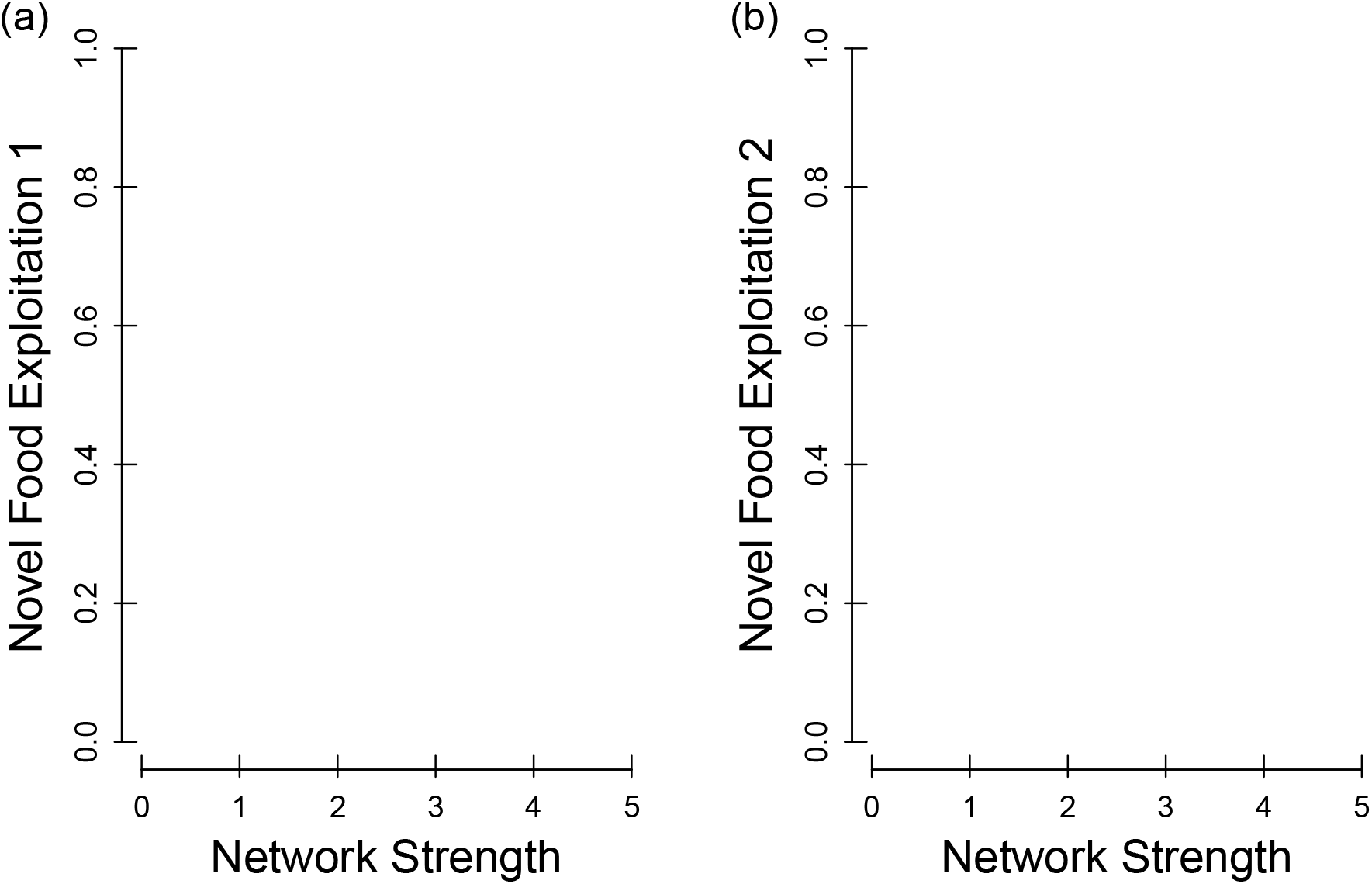
Social centrality novel food exploitation after first use. Prior social centrality (network strength – x axis) and subsequent novel food exploitation (proportion of novel food usage – y axis) after they had overcome any neophobia for the (a) first trial, and (b) second trial. The point positions show the individual data points, point colour shows the colour of the novel food (red or green dyed peanut), point shape shows which experimental site the individual was at (site 1 or site 2), and point size indicates weight of the data point i.e. the total number of detections (at both the novel, and familiar food feeder). The lines show the GLM fit, and the surrounding polygons show the associated standard error around this estimate, with the red lines showing the fit for the red novel food site, the green line showing the fit for the green novel food site, and the black line denoting the overall fit. See Table S10 for full model details.

## DISCUSSION

By quantifying wild great tit social networks, and tracking their behaviour in experimental trials aimed at testing dietary decisions, we found that individuals’ prior social network centrality predicted their subsequent propensity to use novel food, and that this was unrelated to various measures of neophobia. This link between individual sociality (as measured as social network position) and dietary decision making has important implications for understanding how different behavioural strategies influence resource acquisition, and for understanding the emerging consequences of these strategies and decisions ^15^.

Through measuring the sociality of individuals using social network analysis, we were able to quantify the individual level of sociality with this generalisable and powerful approach ^35^. Specifically, we measure individuals’ own social propensity and experienced social environment using metrics that take into account their general sociality (‘strength’ Figure 2), their average bond strength to others (‘edge weight’ Figure S1a;b), and the social centrality of their associates (‘eigenvector centrality’ Figure S1c;d). This approach outperforms simpler methods of attempting to measure sociality ^36^ when making wider inferences e.g. using estimates of group size (Figure S2a;b) or number of group members (Figure S2c;d). Here, the positive relationship between network centrality and novel food usage in this free-living system of socially foraging individuals supports the expectation that competition in social environments can predict foraging strategies in natural populations.

Specifically, individuals that are very social may be able to offset the competitive costs of reduced resources by using other food sources. Furthermore, our approach allowed us to demonstrate that this effect of prior social network centrality on subsequent novel food usage was unlikely to be due to more social birds just generally being more exploratory in this experimental context or less averse to approaching the novel-looking food presented here, as there was no significant relationship between individuals’ social centrality and their delay in approaching the novel food. Thus, it would appear that highly social great tits which may experience a more competitive social environment (i.e., due to having more social ties) may be alleviating the potential costs of competition by foraging more broadly. These findings could be explained by optimal foraging theory ^37–40^ which states that individuals’ foraging decisions should maximise their net rate of energy intake given their environment, as the more socially central great tits (i.e., those potentially experiencing a more competitive social environment) may be more likely to expand their diets by consuming novel food .

Our findings that birds showing the highest degree of dietary conservatism (i.e. those eating least novel food) held the least central network positions, may suggest that these individuals generally experience a reduced level of competition compared to those willing to eat the novel food. In a previous study investigating dietary conservatism and competition in wild-caught captive blue tits (*Cyanistes caeruleus)* ^41^, a high proportion of individuals displayed a strong aversion to the novel food presented to them when foraging alone, i.e. dietary conservatism, preferring to forage only on familiar food. However, with the introduction of a second individual this aversion was quickly overcome, resulting in consumption of novel food. These findings support our own finding which suggest that less social individuals may not experience the same level of competition felt by their more social conspecifics, and thus they may not have to resort to exploiting a novel food source in order to forage efficiently. Furthermore, it could also be argued that these conservative foragers were demonstrating resource partitioning behaviour i.e. reducing dietary overlap with their more social counterparts by excluding this novel food source from their diet ^42^. This kind of behaviour has been demonstrated in other social species, where ‘specialised’ individuals exist. For example, Sheppard, Inger et al. ^16^ found that individual banded mongooses (*Mungos mungo*) with substantially narrower resource niches compared to other members of their social group experienced reduced intraspecific competition through niche partitioning. There are many other instances where individual resource-level specialisation has been examined ^14,15,43^ and where competition has been implicated in driving this resource use variation among individuals of the same population ^44^. Here our results suggest that a potential mechanism regulating these kinds of processes might be linked to social network position. Those more conservatively foraging individuals with fewer foraging social ties, and therefore experiencing a less competitive social environment, do not have to risk expanding their diet to include foods of unknown profitability. In contrast, those with more social ties and therefore more potential competitors can expand their diets and include novel food to mitigate the potential effects of any intraspecific competition for themselves.

The positive effect of social centrality on novel food exploitation may also have consequences for considering the evolution of conspicuous prey defences. Novel conspicuous prey are expected to suffer high initial attack risk from naïve predators ^20,53^ and our results support this idea: Adventurous foragers ‘attacked’ novel food. This makes the initial evolution of conspicuous prey types paradoxical. However, just as birds use social information to find novel food (as specified above), individuals may also transmit social information about prey defences, which can aid the evolution of aposematism ^54,55^ . If more social individuals attack novel prey (as suggested here), they may provide information about prey defences to others, and influence how this social information spreads in the predator population. Further research into the fine-scale interactions between dietary wariness and social learning may be valuable for understanding the evolution of conspicuous prey types.

### Conclusion

By monitoring wild great tit activity at experimental feeders, we showed that socially central individuals are more likely to use novel food than less social individuals. This finding suggests that individuals experiencing a more social associations may be more likely to incorporate novel food resources. Our results also suggest the relationship between social centrality and novel food usage is unlikely to be due to individuals’ initial aversion to first using this new food resource. Therefore, variation in exploratory behaviour in this context, or differences in access to new social information (i.e. variation in discovery rates of the feeders), appear to be improbable drivers of link between sociality and novel food usage. Further expansions of the investigation into social behaviour and individual dietary decisions may now provide insights into topics such as the competition, foraging, sociality, and even the evolution conspicuous prey defences.

## LIMITATIONS OF STUDY

Although the results of the experiment show a clear link between social network centrality and novel food usage, it is also important to highlight limitations of this and areas for future development.

Firstly, while this focus primarily considers competition-driven elements of the relationship between sociality and novel food use, it is important to cosider that social information often shapes foraging decisions. Indeed, information about the profitability and nutritional value of the food may be transmitted to group members through social facilitation ^45^ or local enhancement ^1^. For example, prior work within our great tit population has demonstrated that individuals use social information to locate new foraging locations, and that more central individuals are most likely to learn the location of new resources faster ^46,47^. The relationship between social centrality and information also appears in other species, such as the acquisition of information in social groups of wild baboons (*Papio ursinus*; ^48^. In this study, it is unlikely that social network position shaped the propensity for individual great tits to find the novel food (as it remained in a set location), and also unlikely that it shaped their propensity to try the novel food (as there is no significant relationship between social centrality and timing of using the novel feeder). However, it may be the case that social influence potentially played a role in the extent to which individuals exploited the novel food following discovery of it. For instance, McMahon, Conboy et al. ^49^ showed that conservatively foraging domestic chicks (*Gallus gallus domesticus*) were more willing to consume novel food when they were able to see conspecifics consuming novel food, essentially treating conspecifics as sources of social influence. As such, more socially central great tits may be more likely to be socially associated with others using the novel food resource (simply due to having more social ties) and thus more likely to increase usage of the novel food themselves. On the other hand, less social individuals with fewer social ties may experience less social influence for using the novel food (due to having fewer links to others in general). Indeed, individuals displaying this dietary conservatism may be simply more efficient at exploiting foods with which they are familiar and therefore remain more rigid in their foraging decisions. ^50^. Others have also shown that individuals with higher network centrality may tend to have a more proactive personality ^51^ and that these individuals could also be important in the spread of information because they move more between groups ^52^.

Secondly, the experimental design was limited to two specific colours (red and green) chosen for ‘creating’ the novel food (dyed peanut). Indeed, an additional finding of our experiments was a preference for red novel food over green novel food in this context, as the birds generally preferred the familiar food (standard peanut granules) to a much larger extent when the alternative option was green food compared to when the alternative was red food. Although colour preferences for food are context dependent in birds ^56^, a general preference for red food over green has been reported previously in relation to dietary decision making, such as captive blue tits and great tits preferring red almond flakes over green ^56^, and captive domestic chicks (Gallus gallus domesticus) generally preferring red coloured food over green ^57^ but with other colour preferences varying depending on the types of foods offered ^58^ or experiences prior to being given a colour preference test ^59^.

Addressing these limitations in future research would now be beneficial, particularly in examining how social influence over longer-time periods may govern novel food usage, and by assessing relative novel food preference across a range of contexts and different colours/food types.

## ACKNOWLEDGEMENTS

J.A.F. was supported by a research fellowship from Merton College and BBSRC (BB/S009752/1) and we also acknowledge funding from NERC (NE/S010335/1 and NE/V013483/1) and WildAI (CBR00730). We would also like thank Sam Crafts for his help with additional bird ringing and mist netting prior to the experiments.

## AUTHOR CONTRIBUTIONS

Conceptualisation: KM BCS & JAF. Data Collection and Experimental Design: KM & JAF. Data Analysis: KM & JAF. Results Interpretation: All Authors. Initial Draft Writing: KM & JAF. Revised Draft Writing: All Authors.

## STAR METHODS

### RESOURCE AVAILABILITY

#### Lead Contact

Further information and requests for resources relating to this manuscript should be directed to and will be fulfilled by Keith McMahon

#### Materials availability

This study did not generate new unique reagents.

#### Data and Code availability

Data have been deposited at https://datadryad.org/stash/dataset/doi.10.5061/dryad.3tx95x6fw and are publicly available as of the date of publication. DOIs are listed in the key resources table.

All original code has been deposited at https://zenodo.org/records/10793956 and is publicly available as of the date of publication. DOIs are listed in the key resources table.

Any additional information required to reanalyze the data reported in this paper is available from the lead contact upon request.

## EXPERIMENTAL MODEL AND STUDY PARTICIPANT DETAILS

This study did not use experimental model animals, experimental in-vivo animals, human participants, plants, microbe strains, cell lines, or primary cell cultures.

## METHOD DETAILS

### Study System

Wytham Woods, Oxford, United Kingdom (51° 46′ N, 1° 20′ W) is home to a long-term study population of wild great tits^60^. These birds are captured and tagged with British Trust for Ornithology rings (as adults and as nestlings) during the spring as they breed in the intensively-monitored nest boxes ^60^, and immigrant birds are captured during the winter during regular mist-netting sessions throughout the woodland. As well as recording standard morphological information during capture, since 2007, all captured great tits have also been fitted with radio-frequency identification (RFID) tags. Each RFID tag possesses a unique ID code which allows automated recording of the times and locations of individuals’ occurrence at feeding stations over the winter. Each feeding station consists of a feeding tube with a feeding hole that is equipped with an RFID antenna which successfully records>99% of RFID tagged individuals visits to feeders^61,62^. The feeding stations are set >1m from the ground and surrounded by 1m^3^ wire mesh that protects the equipment from grey squirrels and provides multiple perching points for the birds. These RFID feeding stations allow the recording of individual feeder usage (see Methods: Experiment data) and also the inference of flock structures and arising social networks (see Methods: Social network data). The antennae scan for RFID-tagged individuals 16 times per second from pre-dawn until post-dusk (i.e. over the entirety of the great tits’ foraging hours).

The study was conducted at two separate sites within Wytham Woods, approximately 1km apart, and both sites with similar levels of vegetation cover. Within the timeframe of the study, this 1km distance between sites effectively ensures two separate local populations; of the 105 birds recorded as part of this study, only one individual was observed at both sites (see Supplementary Methods S1). Previous work has estimated that >80% of locally-occurring great tit individuals are RFID tagged51.

### Social Network Data

Prior to beginning the experimental trials, we gathered detailed baseline information regarding individuals’ usage of a familiar food, and their social connections to one another. From 11/01/2018 to 22/01/2018 an RFID feeding station containing non-coloured granulated peanut was placed at each site. Granulated peanut is a familiar food source which is commonly used by great tits in Wytham Woods and the surrounding area, as well as throughout the UK ^63^

Each RFID station automatically recorded the unique identity of each individual detected along with the associated time-stamp. Because these birds forage in loose fission-fusion flocks ^61^, this produced a temporal data stream made up of bursts (as flocks arrive and feed) interspersed with intermittent quiet periods ^64,65^. These bursts of activity (the flocking events) were detected automatically (without the need for subjective specifications) using a Gaussian Mixture Model (GMM – an unsupervised learning algorithm) ^65^ which returns a group-by-individual matrix ^31^ specifying which individuals were detected within each of these flocking events. Following this, social networks can be derived for any desired time period by applying the widely used ‘Simple Ratio Index’ (SRI) ^66^ to the ‘groups’ (i.e. flocking events) observed within that time period, derived as a proportion of flocking events in which the focal dyad (A and B) were seen together as Flocks_A,B_/(Flocks_A_+Flocks_B_-Flocks_A,B_), where Flocks*_A_* is the number of flocks that individual *A* was seen in, irrespective of the observation of *B*. In this way, a weighted, symmetrical, social network was produced for all three periods of the study (baseline, experiment 1, and experiment 2).

In these social networks, the individuals are represented as the network ‘nodes’, and the social connections between them as the network ‘edges’, and the weight of these edges are the dyadic association scores (as specified in the dyadic association matrix). These weights denote the strength of the social affiliation between each of the dyads ^65^.

This approach to calculating social networks has been extensively used for this population and methodological examination of this system has found that the GMM approach outperforms other potential methods of identifying associations ^64,65^. Large-scale observational studies have shown that the derived social networks are consistent across time ^67^ and contexts ^62^, and linked to other processes such as mating ^68,69^, territory acquisition ^62,70^, and information flow ^71^. Furthermore, detailed experimental tests have confirmed the social network’s consistency ^72,73^, and its relation to biologically meaningful outcomes ^74–76^.

We quantified individuals’ social network centrality from the weighted social networks. A common and intuitive metric of social network centrality is weighted strength, which is the sum of the focal individual’s social connections to all other individuals, and is a consistent and repeatable measure of social phenotype in this population ^67^. We also calculated two other measures of social centrality, namely *(i)* ‘average edge weight’ which measures the typical strength of an individual’s social bonds by taking the mean weight of their non-zero dyadic social association scores, and *(ii)* ‘eigenvector centrality’ which measures their position within the wider network by summing the social connections of their associates, and thus represents the sociability of their social associates.

As well as computing these social metrics, we also calculated for each individual the mean size of the flocking events (i.e. the grouping events automatically identified from the feeder co-occurrence records) they occurred in (i.e. their average group size), and the number of unique individuals they were seen with, across all observations. In this way, we were able to separate the influence of individuals’ social network metrics from simpler social measures (see Methods; Statistical Analysis).

### Experiment Data

Each of the experimental trials were carried out after 12 days of baseline data collection. The same general protocol was used at both sites. The first novel food experimental trial took place immediately after the baseline data collection. The single clear-plastic tube RFID feeder (containing familiar food) was swapped for two clear-plastic tube RFID feeders at either side of the original feeder position, within 1m of one another. One of these RFID feeders contained the familiar peanut granules, while the other feeder contained peanut granules which were made novel by dying them either green or red, under standardised methods^41^, for details see supplemental materials section 3. Both feeders were made of transparent plastic to allow the birds to see the colour of the food This experimental trial ran for four days, recording all visits by RFID tagged birds to each of the feeders.

Following this first experimental trial, a second novel food experimental trial was then carried out, in which the feeder containing the novel food was swapped to contain different coloured novel food. In the first experimental trial, the novel food RFID feeder at Site 1 was filled with red-dyed granules while the novel food RFID feeder at Site 2 contained green-dyed granules. In the second experimental trial. This was switched so that the novel food RFID feeder at Site 1 was filled with green-dyed granules while the novel food RFID feeder at Site 2 contained red-dyed granules. In both trials, familiar coloured food was provided in the other feeder at each site. The second experimental trial was carried out for four days (the same length as the first trial), and all visits by RFID tagged birds to the feeders were recorded.

During the experimental trials, we also aimed to reduce any additional influences on the birds’ feeding behaviour that may be caused by either human presence causing disturbance, or through positional effects of feeder placement. We ensured that all required activity at the feeders (i.e. placement changes and associated device checks) were carried out when the great tits were not using the feeders (i.e. after dusk). Even though the familiar-food feeder and the novel food feeder were next to one another (>1m apart), we also aimed to reduce any remaining fine-scale positioning effects by swapping the feeders’ positions every other day during the experiment (see Table S1).

### Quantification and Statistical Analysis

#### Novel Food Usage

For each of the experimental trials, we examined how prior social centrality (i.e. their network centrality before the experiments began) was related to subsequent usage of novel food during the trials. As we aimed to consider individuals’ relative use of the novel food, rather than just their total feeder use in general, we treated the proportion of their total activity which took place on the feeder containing novel food as a measure of individual propensity to use novel food. Therefore, we carried out logistic regressions for each of the trials separately, whereby the response variable in the generalised linear model (GLMs) was set as a binomial variable with the number of detections on the novel food feeder as ‘successes’ and the number of detections on the familiar food feeder during the trial as ‘fails’. In this way, the total feeder usage, and also confidence in their propensity (i.e. strength of their bias/preference) to use novel vs familiar food, was considered directly within the response variable. Because GLMs with binomial error-distributions are vulnerable to over-dispersion, we used a quasi-binomial error distribution, which removed this issue of over-dispersion. The models were set to include fixed effects of the factors that could potentially be related to individual novel food usage propensity. We specified the primary explanatory variable of interest as social network centrality (weighted strength) prior to the experimental trial. For each trial, the social centrality used for the analysis were derived from the period immediately before the trial. As such, for the model assessing the novel food usage during the first trial, individual social centrality calculated from the network directly before the trial began (i.e. during the baseline data collection period) was used. For the model assessing the second trial, weighted strength during the period directly before the second trial began (i.e. during the first trial data collection period) was used. We also aimed to account for other variables that may affect novel food usage, and included site (i.e. which of the two areas the individual was detected in), sex (whether they were male or female), age (specified as either adult, or juvenile), and immigratory status (whether they had arrived in the Wytham Woods study area that year or not) as explanatory variables in the model. In order to directly consider individual differences in feeder usage, we also included the number of detections on the feeders in the period prior to the experimental trial.

Although social centrality was set as weighted network strength in the main models, we also quantified it using other commonly used network metrics (see Methods *Social Network Data*). Therefore, we ran supplementary models using other common measures of social network centrality (average edge weight, eigenvector centrality) calculated from the period prior to the trial, while the rest of the model structure remained the same. Furthermore, it is also possible that other more basic measures of sociality (i.e. non-network based metrics) might act as potential explanatory variables (see Methods *Social Network Data*). To test this, we ran the same models again using each of the simple individual-level social metrics (i.e. not based on networks) obtained from the period prior to the experimental trial (average size of the flocking events they were observed in, and number of unique individuals they were seen with).

#### Neophobia

Individual variation in the observed usage of novel food in the experimental trials could potentially be due to differences in the propensity to first approach the novel food (i.e. avoidance/neophobia) rather than variation in propensity to use the food once any potential neophobia is overcome. We considered this directly by employing the same models as described above, but instead of setting proportional novel food usage as the response variable, we used a binary variable of whether or not they were detected on the feeding perch of the novel food feeding station when they first arrived at the feeding site during the experimental trial. We used a GLM with a quasi-binomial error distribution, and fitted the same fixed effects of the main models (prior social centrality, individual sex, age, immigrant status, experimental site, and prior number of feeding detections).

Another measure of neophobia is the latency to first approach the novel food (as opposed to the ‘likelihood of using the novel food upon the first visit’ as described above). Therefore, we also calculated two related temporal measures of individual novel food neophobia; ‘time to use the novel food since the experiment began’, and ‘time to use the novel food since the individual was first detected during the experimental trial (i.e. time since they first landed on either feeder during the experiment) ’. We set each of these in turn as response variables in the same model structure as described above, but using a gaussian-error distribution instead of binomial due to the distribution of these response variables.

After modelling how the explanatory variables were related to measures of novel food neophobia, we also re-assessed the models examining novel food usage propensity but only considering individuals’ behaviour once any neophobia had been overcome i.e. once the individual had already approached and used the novel-food feeder; Specifically, we re-calculated each individual’s proportional usage of novel and familiar foods but this time only within the time-period following their first detection on the perch of the novel food feeder. Following the primary model structure, we fitted this as the response variable in a GLM with binomial error-structure (with the novel food usage as ‘successes’ and familiar food usage as ‘fails’), along with the explanatory variables (as stated previously) to examine how this predicted novel food usage once individuals had already used the novel food feeder. We additionally evaluated how model structure related to the observed results using randomisations (see below).

#### Network Randomisations

Individuals’ positions within social networks are dependent on one another ^31^. Social network data, by definition, violates the assumption of the independence of data points made under the standard maximum likelihood statistical tests. Therefore, network randomisations are commonly used when estimating the statistical significance of observed parameters computed from standard tests ^31^. Such randomisation techniques allow the creation of null models using a given permutation procedure, and from these null models the same parameters can be re-calculated using the permuted data (instead of the observed data) to provide the distribution of this parameter that is expected given the underlying network structure, and the non-independence of data. More broadly, null models based on permutations of the observed data can also act as an additional, and intuitive, test of significance of observed statistics across various contexts. We employed a hierarchical node attribute permutation procedure controlling for space and time ^62^ whereby individuals were randomly reassigned the attributes (response variable of consideration) of another node individual in the same area during the same period of consideration as themselves. Following this, we re-ran the models and stored the estimated effect size (Coefficient) of each of the predictor variables on the permuted response variable, while keeping everything else in the model the same (i.e. maintaining the exact distributions of all the variables, and the covariance between the predictor variables). By running 10,000 of these permutations, we generated the null distribution of the effect size parameter for each model’s predictor variables and calculated the significance of the observed data test statistics by comparing it to these null distributions. In this way, the p-value (p_rand_) represents each observed statistic’s position within the corresponding null distribution, whereby p_rand_<0.05 indicates that the observed statistic lays outside of the 95% range of the null distribution for this predictor variable (i.e. below the bottom 2.5% or above the top 97.5%, i.e. it detects a significant effect).

## SUPPLEMENTARY INFORMATION CONTENT

### (1) Supplementary Figures

**Figure S1:**
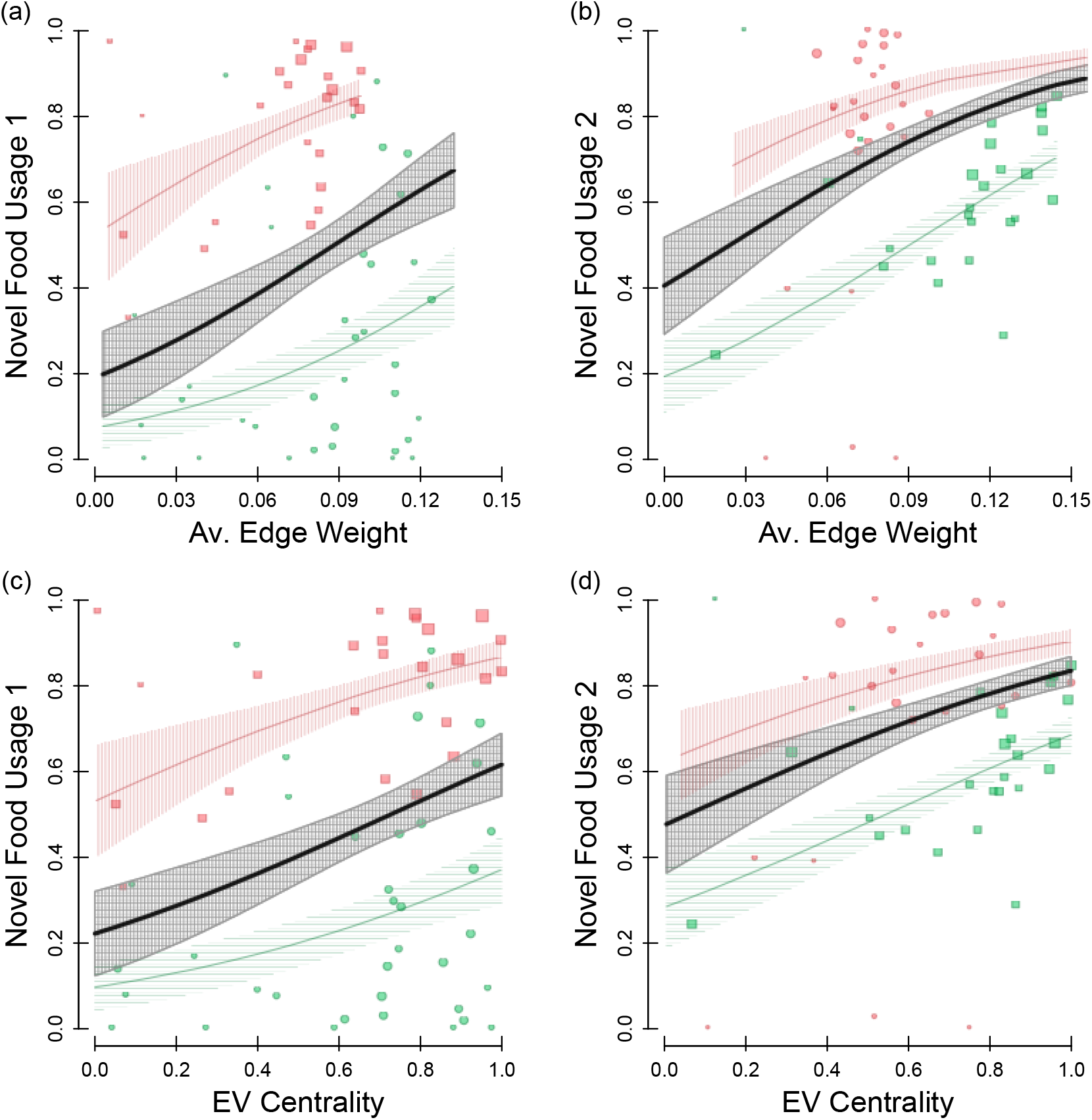
Social centrality metrics and novel food usage, related to Figure 2. Prior social centrality (x axis), as measured as (a-b) Average edge weight and (c-d) eigenvector centrality, and subsequent novel food usage (proportion of novel food usage – y axis) for the (a,c) first trial, and (b,d) second trial. The point positions show the individual data points, point colour shows the colour of the novel food (red or green dyed peanut), point shape shows which experimental site the individual was at (site 1 or site 2), and point size indicates weight of the data point i.e. the total number of detections (at both the novel, and familiar food feeder). The lines show the GLM fit, and the surrounding polygons show the associated standard error around this estimate, with the red lines showing the fit for the red novel food site, the green line showing the fit for the green novel food site, and the black line denoting the overall fit. See Table S3 & S4 for full model details.

**Figure S2:**
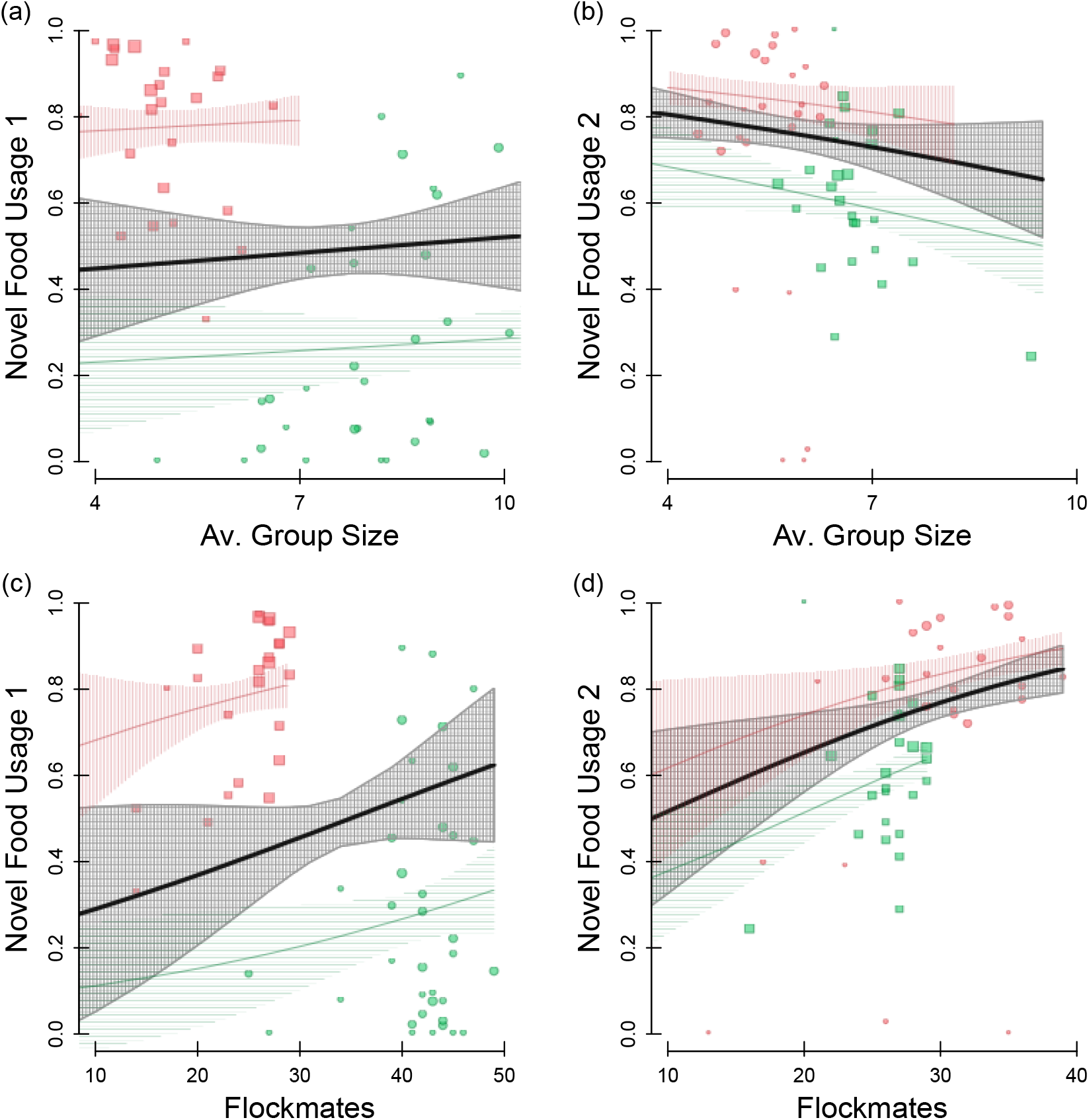
Basic social measures and novel food usage, related to Figure 2. Average group size is the average size of the flocking event that the individual was observed in, and flockmates is the total number of unique individuals the individual was observed occurring with in at least one flocking event. Prior basic measures (x axis), as measured as (a-b) Average group size and (c-d) number of flockmates, and subsequent novel food usage (proportion of novel food usage – y axis) for the (a,c) first trial, and (b,d) second trial. The point positions show the individual data points, point colour shows the colour of the novel food (red or green dyed peanut), point shape shows which experimental site the individual was at (site 1 or site 2), and point size indicates weight of the data point i.e. the total number of detections (at both the novel, and familiar food feeder). The lines show the GLM fit, and the surrounding polygons show the associated standard error around this estimate, with the red lines showing the fit for the red novel food site, the green line showing the fit for the green novel food site, and the black line denoting the overall fit. See Table S5 & S6 for full model details.

### (2) Supplementary Tables

**Table S1:** Summary of experimental procedure, related to Figure 1. The study protocol at each of the sites, showing the phase of the study and food-types used over the data-collection days and the fine-scaling positioning of the feeders within the feeding sites.

| Site | Phase | Day | Food Type | Position |
| --- | --- | --- | --- | --- |
| 1 | Baseline | 1-12 | Familiar | Mid |
|  | Trial 1 | 13-14 | Familiar | Side 1 |
|  |  |  | Green | Side 2 |
|  |  | 15-16 | Familiar | Side 2 |
|  |  |  | Green | Side 1 |
|  | Trial 2 | 16-17 | Familiar | Side 2 |
|  |  |  | Red | Side 1 |
|  |  | 18-19 | Familiar | Side 1 |
|  |  |  | Red | Side 2 |
| 2 | Baseline | 1-12 | Familiar | Mid |
|  | Trial 1 | 13-14 | Familiar | Side 2 |
|  |  |  | Red | Side 1 |
|  |  | 15-16 | Familiar | Side 1 |
|  |  |  | Red | Side 2 |
|  | Trial 2 | 16-17 | Familiar | Side 2 |
|  |  |  | Green | Side 1 |
|  |  | 18-19 | Familiar | Side 1 |
|  |  |  | Green | Side 2 |

**Table S2:**
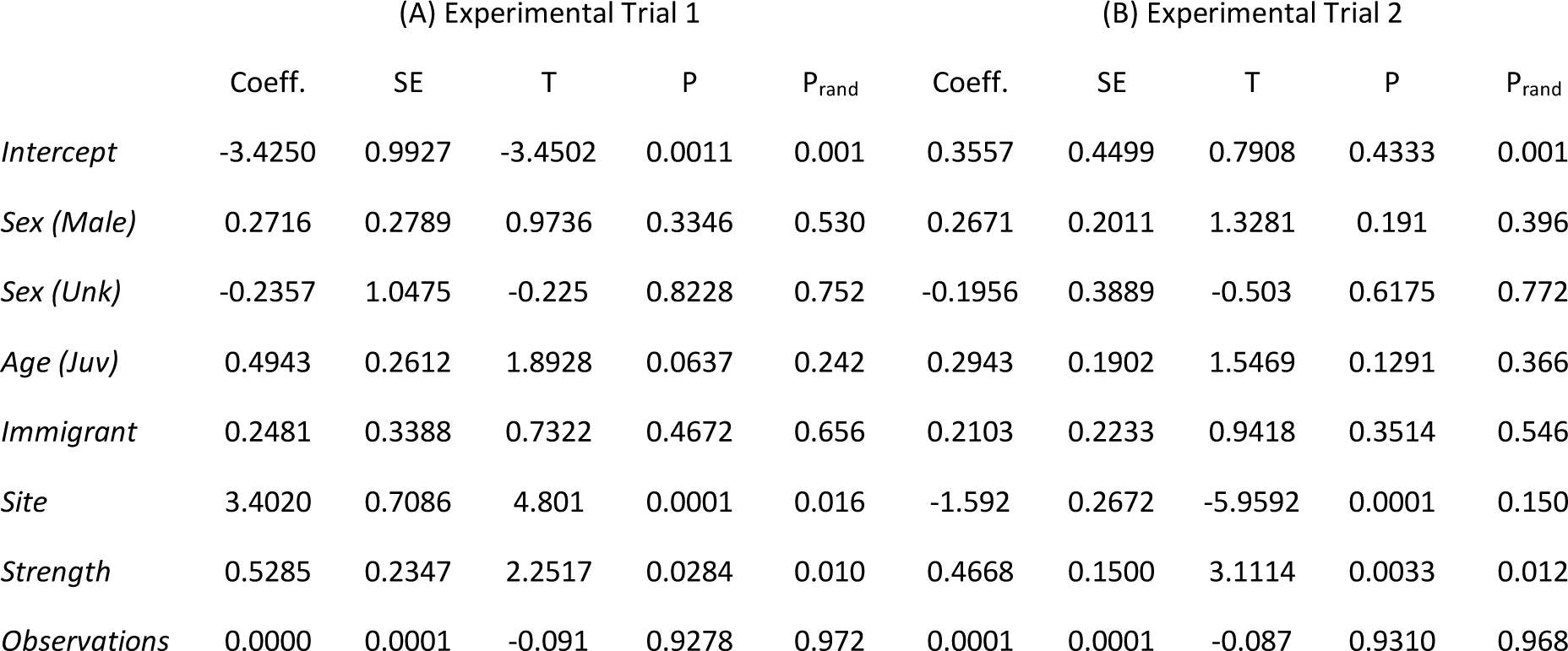
Social network strength and novel food usage model outputs, related to Figure 2. Output of GLMs assessing the relationship between individuals’ propensity to use novel food (response variable) and individuals’ prior network strength (Figure 2 - Main Text), along with the other fitted explanatory variables. Each column holds the test statistics for (A) experimental trial 1 and (B) experimental trial 2. Each row gives the result for each explanatory variable, with ‘Sex’ in relation to female birds, Age in relation to adult birds, Immigrant status in relation to residents, and the ‘Strength’ as weighted network degree directly prior to each experimental trial (see Methods) and ‘Observations’ as the number of records.

**Table S3:** Average edge weight and novel food usage model outputs, related to Figure 2 and Figure S1. Output of GLMs assessing the relationship between individuals’ propensity to use novel food (response variable) and individuals’ average edge weight (Figure S1a;S1b), along with the other fitted explanatory variables. Each column holds the test statistics for (A) experimental trial 1 and (B) experimental trial 2. Each row gives the result for each explanatory variable, with ‘Sex’ in relation to female birds, Age in relation to adult birds, Immigrant status in relation to residents, and the ‘Edge’ as average non-zero edge weight directly prior to each experimental trial (see Methods) and ‘Observations’ as the number of records.

|  | (A) Experimental Trial 1 |  |  |  |  | (B) Experimental Trial 2 |  |  |  |  |
| --- | --- | --- | --- | --- | --- | --- | --- | --- | --- | --- |
|  | Coeff. | SE | T | P | P<sub>rand</sub> | Coeff. | SE | T | P | P<sub>rand</sub> |
| *Intercept* | -2.8021 | 0.7216 | -3.8833 | 0.0001 | 0.0001 | 0.2123 | 0.4475 | 0.4745 | 0.6375 | 0.002 |
| *Sex (Male)* | 0.2257 | 0.2763 | 0.817 | 0.4175 | 0.642 | 0.2974 | 0.1978 | 1.5033 | 0.1399 | 0.348 |
| *Sex (Unk)* | -0.446 | 1.0028 | -0.4447 | 0.6583 | 0.532 | -0.0812 | 0.3914 | -0.2074 | 0.8367 | 0.922 |
| *Age (Juv)* | 0.4891 | 0.261 | 1.8738 | 0.0664 | 0.25 | 0.3282 | 0.1876 | 1.7499 | 0.0871 | 0.302 |
| *Immigrant* | 0.1774 | 0.3361 | 0.5279 | 0.5997 | 0.754 | 0.2717 | 0.2216 | 1.2261 | 0.2267 | 0.436 |
| *Site* | 2.55 | 0.3977 | 6.4116 | 0.0001 | 0.236 | -1.8735 | 0.2753 | -6.8054 | 0.0001 | 0.006 |
| *Edge* | 16.4625 | 7.2095 | 2.2835 | 0.0264 | 0.022 | 15.9091 | 4.5385 | 3.5053 | 0.0011 | 0.004 |
| *Observations* | 0.0001 | 0.0001 | 0.18 | 0.8578 | 0.91 | 0.0001 | 1E-04 | -0.2065 | 0.8374 | 0.928 |

**Table S4:** Eigenvector centrality and novel food usage model outputs, related to Figure 2 and Figure S1. Output of GLMs assessing the relationship between individuals’ propensity to use novel food (response variable) and individuals’ eigenvector centrality (Figure S1c;S1d), along with the other fitted explanatory variables. Each column holds the test statistics for (A) experimental trial 1 and (B) experimental trial 2. Each row gives the result for each explanatory variable, with ‘Sex’ in relation to female birds, Age in relation to adult birds, Immigrant status in relation to residents, and the ‘Eigenvector’ as weighted eigenvector centrality directly prior to each experimental trial (see Methods) and ‘Observations’ as the number of records.

|  | (A) Experimental Trial 1 |  |  |  |  | (B) Experimental Trial 2 |  |  |  |  |
| --- | --- | --- | --- | --- | --- | --- | --- | --- | --- | --- |
|  | Coeff. | SE | T | P | P<sub>rand</sub> | Coeff. | SE | T | P | P<sub>rand</sub> |
| *Intercept* | -2.4432 | 0.5825 | -4.1941 | 0.0001 | 0.001 | 0.3543 | 0.4713 | 0.7517 | 0.4562 | 0.001 |
| *Sex (Male)* | 0.2578 | 0.2787 | 0.9249 | 0.3591 | 0.554 | 0.2684 | 0.2051 | 1.3089 | 0.1973 | 0.396 |
| *Sex (Unk)* | -0.3282 | 1.0023 | -0.3274 | 0.7446 | 0.652 | -0.2009 | 0.3947 | -0.5091 | 0.6132 | 0.758 |
| *Age (Juv)* | 0.5059 | 0.2616 | 1.934 | 0.0584 | 0.226 | 0.2866 | 0.1923 | 1.4901 | 0.1433 | 0.384 |
| *Immigrant* | 0.2423 | 0.3395 | 0.7138 | 0.4784 | 0.656 | 0.2187 | 0.2278 | 0.9598 | 0.3424 | 0.538 |
| *Site* | 2.3106 | 0.3414 | 6.7671 | 0.0001 | 0.718 | -1.4989 | 0.2714 | -5.5235 | 0.0001 | 0.302 |
| *Eigenvector* | 1.7357 | 0.7585 | 2.2881 | 0.0261 | 0.05 | 1.7099 | 0.5834 | 2.9307 | 0.0053 | 0.012 |
| *Observations* | 0.0000 | 0.0001 | -0.3856 | 0.7013 | 0.818 | 0.0000 | 1E-04 | -0.0523 | 0.9585 | 0.988 |

**Table S5:** Mean gathering event size and novel food usage model outputs, related to Figure 2 and Figure S2. Output of GLMs assessing the relationship between individuals’ propensity to use novel food (response variable) and individuals’ average flock size (Figure S2a;S2b), along with the other fitted explanatory variables. Each column holds the test statistics for (A) experimental trial 1 and (B) experimental trial 2. Each row gives the result for each explanatory variable, with ‘Sex’ in relation to female birds, Age in relation to adult birds, Immigrant status in relation to residents, and the ‘Flock size’ as mean number of individuals within each flocking event the individual was observed in directly prior to each experimental trial (see Methods) and ‘Observations’ as the number of records.

|  | (A) Experimental Trial 1 |  |  |  |  | (B) Experimental Trial 2 |  |  |  |  |
| --- | --- | --- | --- | --- | --- | --- | --- | --- | --- | --- |
|  | Coeff. | SE | T | P | P<sub>rand</sub> | Coeff. | SE | T | P | P<sub>rand</sub> |
| *Intercept* | -1.7901 | 1.6104 | -1.1116 | 0.2712 | 0.278 | 2.2743 | 0.8448 | 2.692 | 0.01 | 0.422 |
| *Sex (Male)* | 0.1825 | 0.2921 | 0.6245 | 0.5349 | 0.704 | 0.0399 | 0.2023 | 0.1973 | 0.8445 | 0.904 |
| *Sex (Unk)* | -0.4471 | 0.9903 | -0.4515 | 0.6535 | 0.532 | -0.476 | 0.4422 | -1.0765 | 0.2876 | 0.412 |
| *Age (Juv)* | 0.5458 | 0.2779 | 1.9639 | 0.0547 | 0.202 | 0.1677 | 0.2003 | 0.8371 | 0.4071 | 0.606 |
| *Immigrant* | 0.1904 | 0.3613 | 0.527 | 0.6003 | 0.732 | -0.0225 | 0.2284 | -0.0986 | 0.9219 | 0.930 |
| *Site* | 2.1875 | 0.6794 | 3.2195 | 0.0022 | 0.930 | -1.3842 | 0.3668 | -3.7738 | 0.0001 | 0.534 |
| *Flock Size* | 0.0487 | 0.1684 | 0.2895 | 0.7733 | 0.676 | -0.1404 | 0.1526 | -0.9199 | 0.3627 | 0.232 |
| *Observations* | 0.0001 | 0.0001 | 1.1048 | 0.2741 | 0.322 | 0.0000 | 0.0001 | 2.3104 | 0.0256 | 0.134 |

**Table S6:** Unique flockmates and novel food usage model outputs, related to Figure 2 and Figure S2. Output of GLMs assessing the relationship between individuals’ propensity to use novel food (response variable) and their number of unique flockmates (Figure S2c;S2d), along with the other fitted explanatory variables. Each column holds the test statistics for (A) experimental trial 1 and (B) experimental trial 2. Each row gives the result for each explanatory variable, with ‘Sex’ in relation to female birds, Age in relation to adult birds, Immigrant status in relation to residents, and the ‘Flockmates’ as sum of the number of unique individuals seen in the same flocking events as themselves directly prior to each experimental trial (see Methods) and ‘Observations’ as the number of records.

|  | (A) Experimental Trial 1 |  |  |  |  | (B) Experimental Trial 2 |  |  |  |  |
| --- | --- | --- | --- | --- | --- | --- | --- | --- | --- | --- |
|  | Coeff. | SE | T | P | P<sub>rand</sub> | Coeff. | SE | T | P | P<sub>rand</sub> |
| *Intercept* | -2.7682 | 1.9309 | -1.4337 | 0.1574 | 0.028 | -0.2265 | 1.2268 | -0.1846 | 0.8544 | 0.001 |
| *Sex (Male)* | 0.2176 | 0.2875 | 0.7568 | 0.4525 | 0.628 | 0.1025 | 0.2014 | 0.5089 | 0.6134 | 0.732 |
| *Sex (Unk)* | -0.3795 | 1.0127 | -0.3747 | 0.7093 | 0.596 | -0.4656 | 0.4018 | -1.1587 | 0.2528 | 0.416 |
| *Age (Juv)* | 0.5626 | 0.2698 | 2.0855 | 0.0418 | 0.184 | 0.1811 | 0.1966 | 0.9212 | 0.362 | 0.576 |
| *Immigrant* | 0.3196 | 0.3807 | 0.8396 | 0.4048 | 0.564 | 0.0231 | 0.2252 | 0.1025 | 0.9188 | 0.960 |
| *Site* | 2.6957 | 0.9638 | 2.7968 | 0.0071 | 0.344 | -1.1998 | 0.3843 | -3.1216 | 0.0032 | 0.874 |
| *Flockmates* | 0.0361 | 0.0479 | 0.7535 | 0.4544 | 0.180 | 0.0565 | 0.0385 | 1.4674 | 0.1494 | 0.100 |
| *Observations* | 0.0000 | 0.0001 | 0.3305 | 0.7423 | 0.740 | 0.0000 | 0.0001 | 1.4246 | 0.1613 | 0.326 |

**Table S7:** Social network strength and first feeder used model outputs, related to Figure 3. Output of GLMs assessing the relationship between whether individuals are first detected on the novel food feeder when they first arrive at the experimental trial and their prior network strength (Figure 3 - Main Text), along with the other fitted explanatory variables. Each column holds the test statistics for (A) experimental trial 1 and (B) experimental trial 2. Each row gives the result for each explanatory variable, with ‘Sex’ in relation to female birds, Age in relation to adult birds, Immigrant status in relation to residents, and the ‘Strength’ as weighted network degree directly prior to each experimental trial (see Methods) and ‘Observations’ as the number of records.

|  | (A) Experimental Trial 1 |  |  |  |  | (B) Experimental Trial 2 |  |  |  |  |
| --- | --- | --- | --- | --- | --- | --- | --- | --- | --- | --- |
|  | Coeff. | SE | T | P | P<sub>rand</sub> | Coeff. | SE | T | P | P<sub>rand</sub> |
| *Intercept* | -0.667 | 1.2864 | -0.5185 | 0.6064 | 0.85 | 1.1073 | 1.3439 | 0.824 | 0.4146 | 0.074 |
| *Sex (Male)* | 0.4993 | 0.8285 | 0.6026 | 0.5496 | 0.51 | -0.7047 | 0.7176 | -0.982 | 0.3317 | 0.402 |
| *Sex (Unk)* | 1.2155 | 1.9456 | 0.6247 | 0.535 | 0.378 | -19.37 | 2168.40 | -0.0089 | 0.9929 | 0.006 |
| *Age (Juv)* | -0.0508 | 0.8543 | -0.0595 | 0.9528 | 0.98 | 0.6231 | 0.7775 | 0.8015 | 0.4274 | 0.448 |
| *Immigrant* | 0.892 | 1.1421 | 0.781 | 0.4386 | 0.386 | -1.4231 | 1.1392 | -1.2492 | 0.2185 | 0.196 |
| *Site* | -0.4084 | 1.1975 | -0.341 | 0.7345 | 0.164 | 1.2332 | 0.9898 | 1.246 | 0.2197 | 0.088 |
| *Strength* | 0.2896 | 0.413 | 0.7012 | 0.4865 | 0.29 | -0.5507 | 0.5742 | -0.9592 | 0.3429 | 0.284 |
| *Observations* | 0.0001 | 0.0001 | -1.9471 | 0.0573 | 0.012 | 0.0000 | 0.0001 | -0.6424 | 0.5241 | 0.526 |

**Table S8:** Social network strength and time delay to use novel food, related to Figure 3. Output of LMs assessing the relationship between the amount of time taken for each individual to first land on the feeding perch of the novel food (quantified as time of day they were first recorded on the novel food), and their prior network strength, along with the other fitted explanatory variables. Each column holds the test statistics for (A) experimental trial 1 and (B) experimental trial 2. Each row gives the result for each explanatory variable, with ‘Sex’ in relation to female birds, Age in relation to adult birds, Immigrant status in relation to residents, and the ‘Strength’ as weighted network degree directly prior to each experimental trial (see Methods) and ‘Observations’ as the number of records.

|  | (A) Experimental Trial 1 |  |  |  |  | (B) Experimental Trial 2 |  |  |  |  |
| --- | --- | --- | --- | --- | --- | --- | --- | --- | --- | --- |
|  | Coeff. | SE | T | P | P<sub>rand</sub> | Coeff. | SE | T | P | P<sub>rand</sub> |
| *Intercept* | 38898 | 2064 | 18.84 | 0.0001 | 0.334 | 38958 | 3802 | 10.25 | 0.0001 | 0.614 |
| *Sex (Male)* | 300 | 1125 | 0.2668 | 0.7908 | 0.784 | 1606 | 2079 | 0.7727 | 0.4442 | 0.468 |
| *Sex (Unk)* | 1439 | 2871 | 0.5011 | 0.6187 | 0.4 | 10075 | 4696 | 2.1456 | 0.0379 | 0.038 |
| *Age (Juv)* | -2302 | 1063 | -2.1646 | 0.0356 | 0.014 | -4876 | 2092 | -2.33 | 0.0248 | 0.028 |
| *Immigrant* | 519 | 1452 | 0.3577 | 0.7222 | 0.62 | 102 | 2753 | 0.037 | 0.9707 | 0.926 |
| *Site* | -2875 | 1696 | -1.6956 | 0.0967 | 0.276 | 1929 | 2575 | 0.7492 | 0.458 | 0.34 |
| *Strength* | -377 | 559 | -0.6739 | 0.5038 | 0.466 | -125 | 1487 | -0.084 | 0.9332 | 0.944 |
| *Observations* | 0.2099 | 0.2216 | 0.9472 | 0.3485 | 0.388 | -1.5294 | 1.4881 | -1.028 | 0.3101 | 0.458 |

**Table S9:**
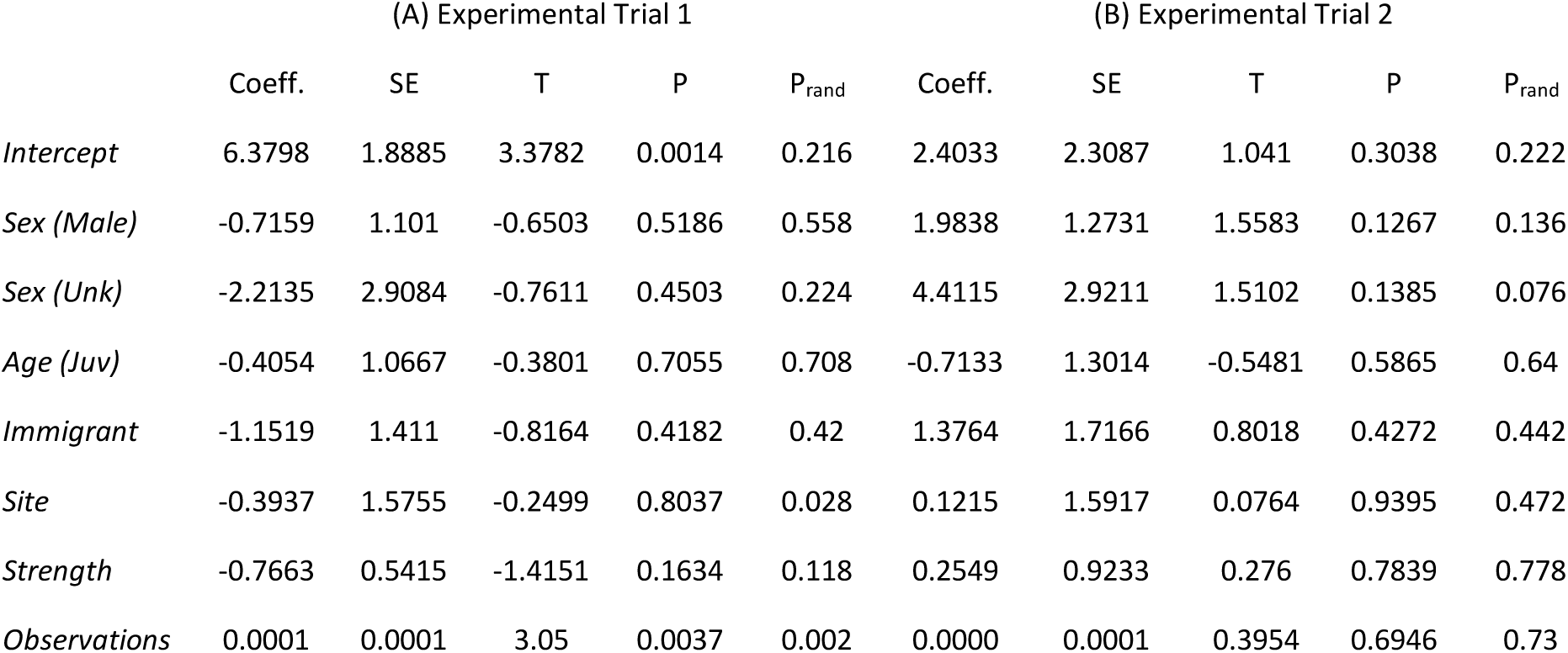
Social network strength and overall time delay to use novel food, related to Figure 3. Output of LMs assessing the relationship between the amount of time taken for each individual to first land on the feeding perch of the novel food (quantified as total elapsed foraging time since they were first detected at the site during the trial – log transformed), and their prior network strength, along with the other fitted explanatory variables. Each column holds the test statistics for (A) experimental trial 1 and (B) experimental trial 2. Each row gives the result for each explanatory variable, with ‘Sex’ in relation to female birds, Age in relation to adult birds, Immigrant status in relation to residents, and the ‘Strength’ as weighted network degree directly prior to each experimental trial (see Methods) and ‘Observations’ as the number of records.

|  | (A) Experimental Trial 1 |  |  |  |  | (B) Experimental Trial 2 |  |  |  |  |
| --- | --- | --- | --- | --- | --- | --- | --- | --- | --- | --- |
|  | Coeff. | SE | T | P | P<sub>rand</sub> | Coeff. | SE | T | P | P<sub>rand</sub> |
| *Intercept* | 6.3798 | 1.8885 | 3.3782 | 0.0014 | 0.216 | 2.4033 | 2.3087 | 1.041 | 0.3038 | 0.222 |
| *Sex (Male)* | -0.7159 | 1.101 | -0.6503 | 0.5186 | 0.558 | 1.9838 | 1.2731 | 1.5583 | 0.1267 | 0.136 |
| *Sex (Unk)* | -2.2135 | 2.9084 | -0.7611 | 0.4503 | 0.224 | 4.4115 | 2.9211 | 1.5102 | 0.1385 | 0.076 |
| *Age (Juv)* | -0.4054 | 1.0667 | -0.3801 | 0.7055 | 0.708 | -0.7133 | 1.3014 | -0.5481 | 0.5865 | 0.64 |
| *Immigrant* | -1.1519 | 1.411 | -0.8164 | 0.4182 | 0.42 | 1.3764 | 1.7166 | 0.8018 | 0.4272 | 0.442 |
| *Site* | -0.3937 | 1.5755 | -0.2499 | 0.8037 | 0.028 | 0.1215 | 1.5917 | 0.0764 | 0.9395 | 0.472 |
| *Strength* | -0.7663 | 0.5415 | -1.4151 | 0.1634 | 0.118 | 0.2549 | 0.9233 | 0.276 | 0.7839 | 0.778 |
| *Observations* | 0.0001 | 0.0001 | 3.05 | 0.0037 | 0.002 | 0.0000 | 0.0001 | 0.3954 | 0.6946 | 0.73 |

**Table S10:** Social network strength and novel food exploitation after first use, related to Figure 4. Output of GLMs assessing the relationship between individuals’ propensity to use novel food after they had already first tried the novel food feeder (response variable) and individuals’ prior network strength (Figure 4 - Main Text), along with the other fitted explanatory variables. Each column holds the test statistics for (A) experimental trial 1 and (B) experimental trial 2. Each row gives the result for each explanatory variable, with ‘Sex’ in relation to female birds, Age in relation to adult birds, Immigrant status in relation to residents, and the ‘Strength’ as weighted network degree directly prior to each experimental trial (see Methods) and ‘Observations’ as the number of records.

|  | (A) Experimental Trial 1 |  |  |  |  | (B) Experimental Trial 2 |  |  |  |  |
| --- | --- | --- | --- | --- | --- | --- | --- | --- | --- | --- |
|  | Coeff. | SE | T | P | P<sub>rand</sub> | Coeff. | SE | T | P | P<sub>rand</sub> |
| *Intercept* | -3.5136 | 1.0769 | -3.2627 | 0.002 | 0.001 | 0.4391 | 0.5057 | 0.8682 | 0.3902 | 0.001 |
| *Sex (Male)* | 0.316 | 0.3108 | 1.0167 | 0.3143 | 0.508 | 0.4231 | 0.2235 | 1.8928 | 0.0653 | 0.210 |
| *Sex (Unk)* | -0.1159 | 1.2082 | -0.0959 | 0.924 | 0.886 | -0.0665 | 0.4319 | -0.1539 | 0.8784 | 0.948 |
| *Age (Juv)* | 0.5041 | 0.2892 | 1.7429 | 0.0876 | 0.264 | 0.3009 | 0.2112 | 1.4244 | 0.1617 | 0.386 |
| *Immigrant* | 0.2262 | 0.3783 | 0.598 | 0.5526 | 0.682 | 0.3363 | 0.2452 | 1.3718 | 0.1774 | 0.364 |
| *Site* | 3.6381 | 0.7803 | 4.6627 | 0.0001 | 0.014 | -1.8279 | 0.3112 | -5.8741 | 0.0001 | 0.158 |
| *Strength* | 0.5483 | 0.2551 | 2.1492 | 0.0366 | 0.006 | 0.5324 | 0.1677 | 3.1742 | 0.0028 | 0.010 |
| *Observations* | 0.0001 | 0.0001 | 0.0207 | 0.9836 | 0.994 | 0.0001 | 1E-04 | -0.4635 | 0.6454 | 0.802 |

### (3) Supplementary Methods S1 Procedure of dyeing peanut granules, related to STAR METHODS

The green dye for the food was prepared by mixing O’Brien’s (Citywest, Dublin 24, Ireland) liquid green 90 food colouring in the ratio of 5 ml dye to 500 ml water. This solution was then mixed with 500 g of kibbled peanut. The mixture was placed in an oven at 50°C for 20–30 min until dry. This was repeated with O’Brien’s Christmas Red for the red dyed peanut.

